# Ultra-fast electroporation of giant unilamellar vesicles — Experimental validation of a molecular model

**DOI:** 10.1101/2020.01.01.890137

**Authors:** Esin B. Sözer, Sourav Haldar, Paul S. Blank, Federica Castellani, P. Thomas Vernier, Joshua Zimmerberg

**Affiliations:** Frank Reidy Research Center for Bioelectrics, Old Dominion University, Norfolk, VA 23508, USA; Section on Integrative Biophysics, *Eunice Kennedy Shriver* National Institute of Child Health and Human Development, Bethesda, MD 20892, USA; Department of Cell and Molecular Biology, Molecular Biophysics, Uppsala University, Sweden; Biomedical Engineering Institute, Frank Batten College of Engineering and Technology Old Dominion University, Norfolk, VA 23529, USA

## Abstract

Delivery of molecules to cells via electropermeabilization (electroporation) is a common procedure in laboratories and clinics. However, despite a long history of theoretical effort, electroporation protocols are still based on trial and error because the biomolecular structures and mechanisms underlying this phenomenon have not been established. Electroporation models, developed to explain observations of electrical breakdown of lipid membranes, describe the electric field-driven formation of pores in lipid bilayers. These transient pore models are consistent with molecular dynamics simulations, where field-stabilized lipid pores form within a few nanoseconds and collapse within tens of nanoseconds after the field is removed. Here we experimentally validate this nanoscale restructuring of bio-membranes by measuring the kinetics of transport of the impermeant fluorescent dye calcein into lipid vesicles exposed to ultrashort electric fields (6 ns and 2 ns), and by comparing these results to molecular simulations. Molecular transport after vesicle permeabilization induced by multiple pulses is additive for interpulse intervals as short as 50 ns, while the additive property of transport is no longer observed when the interval is reduced to 0 ns, consistent with the lifetimes of lipid electropores in molecular simulations. These results show that lipid vesicle responses to pulsed electric fields are significantly different from those of living cells where, for similar pulse properties, the uptake of fluorescent dye continues for several minutes.

**Graphical abstract:** 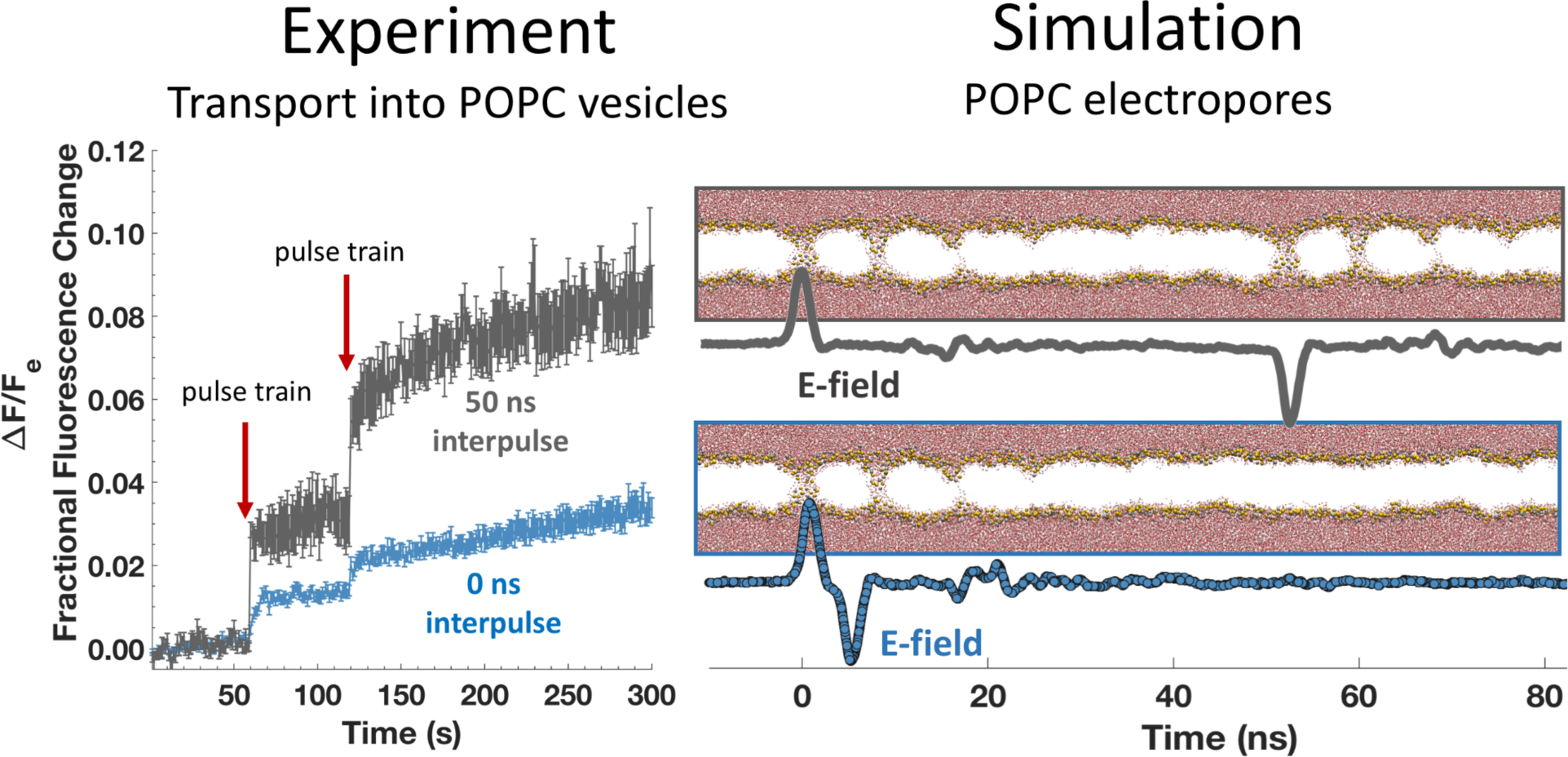

## Main text

Reversible electropermeabilization (electroporation) is widely used in gene and drug delivery, gene editing, and electrofusion, but the structures and mechanisms associated with the electrical disruption of biological membranes have not been established conclusively, despite decades of study^1–7^. Early models of pore formation, based on the interplay of surface and line tensions around a membrane opening^8–9^ were validated using planar lipid bilayer conductance data.^3, 10, 11^ In this theoretical framework, a transmembrane potential (*V_m_*) lowers the energy (*W*) required for the formation of a hydrophilic pore of radius *r*:

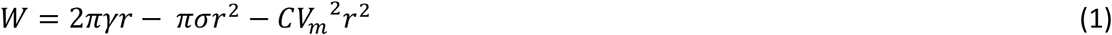

where *γ* is the line tension, *σ* is the surface tension, and *C* is a constant.^3, 8, 9^ This formalism was applied to descriptions of electroporation-based transport of molecules into cells.^12–14^

Molecular dynamics (MD) simulations provide physics-anchored reference points for the calculated behavior of electrically stressed lipid bilayers (the simplest biological membranes). MD simulations show how applied electric fields stabilize random incursions of water into the membrane interior and how lipid pores form within nanoseconds as phospholipids reorganize around these water bridges.^15–18^ Annihilation of lipid electropores takes 10–100 times longer than pore creation.^17^ Pore creation and annihilation times, arising from the molecular model can, in principle, be validated with experiments with simple membrane systems.

Most reports of nanosecond pulsed electric field effects on biomembranes are based on permeabilization of cells or tissues, which includes cell membrane and cellular physiological complexity.^19–29^ A few studies have explored nanosecond pulsed electric field-induced transport across artificial membranes or lipid vesicles,^30–32^ but none has measured transport kinetics after nanosecond pulse permeabilization of lipid membranes.^33^

Here we connect pore creation and annihilation times from MD simulations of phospholipid bilayers with experimental observations of transport kinetics into giant unilamellar vesicles (GUVs) exposed to ultrashort electric pulses (2 ns and 6 ns). We show that a large fraction of the membrane-permeabilizing pores formed under these conditions opens and closes within nanoseconds. We compare these experimental results to those obtained on cells when appropriate.^19–29^ GUVs were made using the gel-assisted formation method,^34–36^ which has several advantages over the electro-formation methods^35^ commonly used in experimental studies of GUVs in pulsed electric fields.^31, 37–39^ For fluorescence imaging simultaneous with electrical stimulation, labeled vesicles were positioned between parallel wire electrodes^22^ in the field of view. Calcein was chosen as a small, membrane-impermeant, fluorescent molecule that does not interact with lipids during transport across membranes due to its hydrophilic properties.^40–42^ Figure 1a shows a typical confocal image of GUVs.

**Figure 1.**
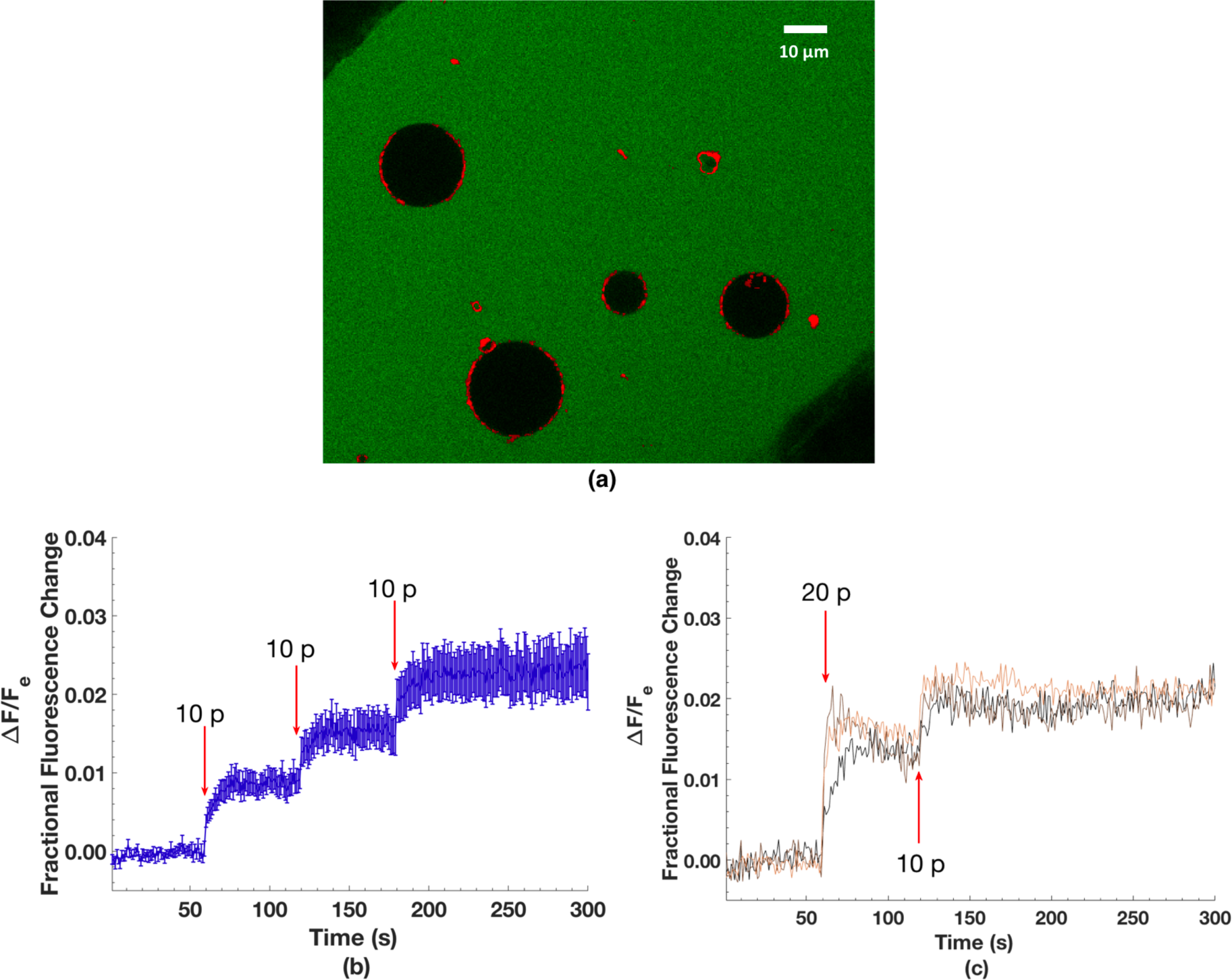
(a) Confocal image of POPC GUVs stained with DiD in 200 µM calcein solution at the bottom of a cover glass chamber and between tungsten wire electrodes. Green represents calcein fluorescence, red DiD. (b) Intravesicular calcein fluorescence of GUVs exposed to three 1 kHz trains of ten 6 ns electric pulses (at field strengths of 35 MV/m) delivered 1, 2, and 3 minutes into the recording (n = 13). Arrows indicate the time of each train of pulses. Error bars are standard error of the mean. (c). Individual fractional fluorescence change traces of three GUVs first exposed to two 1 kHz trains of ten pulses separated by 500 ms (total of 20 pulses, 20 p) and then exposed to one train of ten pulses (10 p). In both the 20 p and 10 p trains, each pulse was 6 ns in duration and 35 MV/m in field strength, and delivered one and two minutes into the recording respectively. One frame per second recordings.

Exposure of 10 ms of a 1 kHz train of electrical pulses (each pulse of duration 6 ns, with a field of 35 MV/m) results in a stepwise increase of intravesicular calcein fluorescence (Fig. 1b), with a faster rise time than that observed with cells after similar exposures.^22^ The increase in fluorescence is proportional to the number of pulses and is additive for successive exposures (Fig. 1c), consistent with the hypothesis that electric fields induce opening of lipid pores and calcein transport through the pores, most or all of which close during the 1 ms interpulse interval. The intravesicular calcein concentration is approximately 1% of the extracellular concentration for 10 pulses and 2% for 20 pulses – 2 µM and 4 µM, respectively. This degree of entry can be described by simple electrodiffusion through a small pore (Supplementary Material). The correlation between GUV size and molecular transport is not positive in contrast with experiments using long pulse durations (100 µs and 5 ms)^39^, which is consistent with the membrane charging time constant for GUVs (~100 ns) being much longer than the duration of these pulses (Figure S2, Supplementary Material).

Next, GUV permeabilization was tested for sensitivity to direction and sequence of electric field i.e. exposure to unipolar and bipolar nanosecond pulses. In cells, the transport of small molecules caused by a unipolar pulse can be attenuated or cancelled by a closely following pulse of opposite polarity.^28,29,42–47^ In other words, a unipolar pulse is more effective in causing small molecule transport into cells than a bipolar pulse of double the duration. 2 ns unipolar pulses and bipolar pulses applied to GUV with no interpulse delay (negative pulse immediately follows the positive pulse without any intervening interval of time) results in equal transport of calcein, whereas a short delay of 50 ns between the bipolar pulse phases results in twice the transport (Fig. 2). Stated another way, for a bipolar pulse with no delay between phases (5 ns interval between the peaks), the negative phase does not affect the permeabilization caused by the preceding positive phase. Note that if GUV membrane conductivity does not change during the 2 ns of the first bipolar pulse phase, then the second phase of the bipolar pulse should discharge the membrane, resulting in no transport across the membrane, similar to the bipolar cancellation effect observed with cells.^42^

**Figure 2.**
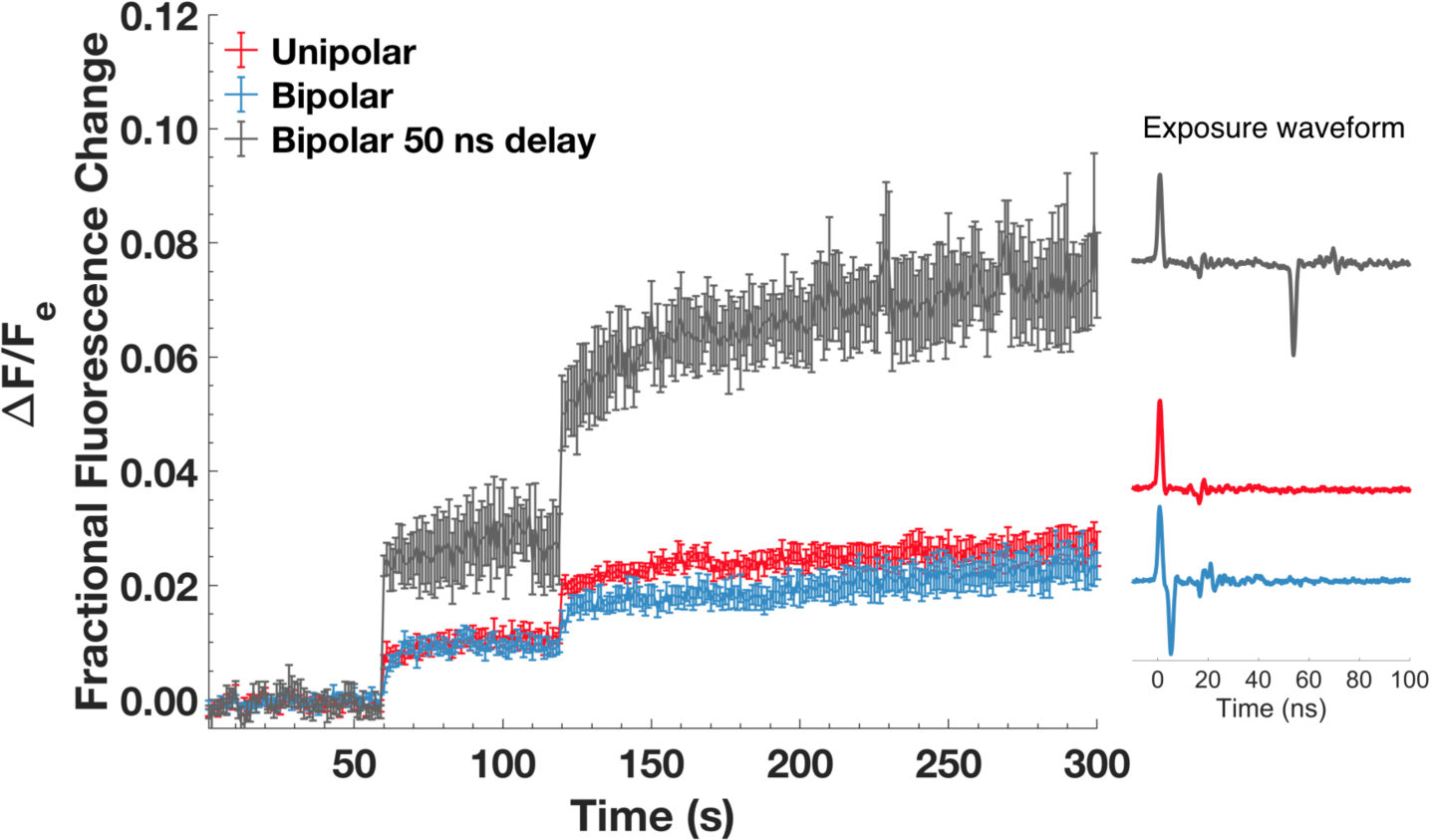
Calcein transport as a result of exposure to unipolar (red, n = 9), bipolar with no interpulse delay (blue, n = 10), or bipolar with 50 ns interpulse delay (gray, n = 4). Trains of 40 pulses (45 MV/m field strength and 2 ns duration) were delivered at a 1 kHz repetition rate 60 and 120 s into the recording. Error bars show standard error of the mean. (Note that the first and second phases of a 2 ns bipolar pulse are each 2 ns in duration.)

The results in Figure 2 can be analyzed within the framework of membrane charging (and discharging) in the standard electroporation models.^3,10,12,14^ We postulate that the positive applied potential of a bipolar pulse permeabilizes the membrane and increases the membrane conductance to a sufficiently high value. Subsequently, when the negative applied potential arrives without any delay, no permeabilizing transmembrane potential develops across the GUV membrane because of its high conductance. Thus, the negative phase of the bipolar pulse cannot significantly change the total transport through existing pores and is not predicted (by these models) to affect membrane permeabilization-related endpoints. Since the second, undelayed phase of the bipolar pulse causes no significant change in total calcein transport, the GUV membrane must be conductive during the second phase of the pulse, and thus most or all of the conductive electropores are formed within the 2 ns duration of the first phase. Even though the total transport is equal, a slower kinetics is observed in bipolar pulse exposures compared to unipolar or bipolar with 50 ns interpulse delay.

Interestingly, the kinetics of the calcein fluorescence increase after 2 ns bipolar pulse exposures (without any interpulse delay) are similar to those observed with 6 ns unipolar pulses (Figure 1b, Figure S3), where a fast transition is followed by a slower increase. The fast process is consistent with the formation of pores observed in molecular dynamics simulations. We hypothesize that the slower process is due to longer lifetime pores, which are more likely to form when the induced membrane potential is sustained longer (i.e. longer pulse durations). Lipid pores with longer lifetimes are observed experimentally and in simulations when a low membrane potential is present.^3,7,48^ This hypothesis also suggests an increase in the number of longer-lasting pores independent of electric field direction, consistent with our molecular dynamics simulations, where field reversal does not affect electropore lifetime. If the second phase of the bipolar pulse is delayed even 50 ns after the first phase of the bipolar pulse, however, the observed additive effect of the second phase on calcein transport is consistent with the charging and permeabilization of a membrane that is no longer significantly conductive – that is, a large fraction of the pore population closes within 50 ns. This roughly 50 ns lifetime of the membrane conductive state is consistent with molecular simulations of lipid electropore annihilation.^17^

Moreover, these data show that significant attenuation or cancellation of molecular transport with electric field reversal, which is observed with live cells,^42^ does not occur in purely lipidic systems. The absence of attenuation in a lipidic system supports the hypothesis that the mechanisms for molecular transport in electropermeabilized living cells involve more than transport through lipidic pores.^22^

To further characterize lipid electropore lifetime after removal of an external electric field, we generated and stabilized pores in molecular simulations of POPC bilayers (128 POPC, ~12000 H_2_O, and 22 K^+^ and 22 Cl^-^, corresponding to 125 mM KCl, close to the KCl concentration in the GUV suspensions). Pores are created by a porating field of 250 MV/m and then stabilized with sustaining fields of 40, 50, and 60 MV/m (corresponding to pore radii of 1.3 nm, 2.0 nm, and 2.4 nm, respectively) for 100 ns,^48^ with three simulations for each condition. The applied field was reduced to zero, and pore radius was monitored for another 100 ns.

Figure 3a shows the evolution of pore size 50 ns before and after removal of the sustaining electric field. Pore radius decreases from a value determined by the sustaining field to about 1 nm, and then remains in this metastable state for 10–35 ns before the pore closes (Fig. 3a). A typical electropore just before complete annihilation is shown in Figure 3b. Pore annihilation time is stochastic and is not correlated with the initial pore size (Fig. 3c).

**Figure 3.**
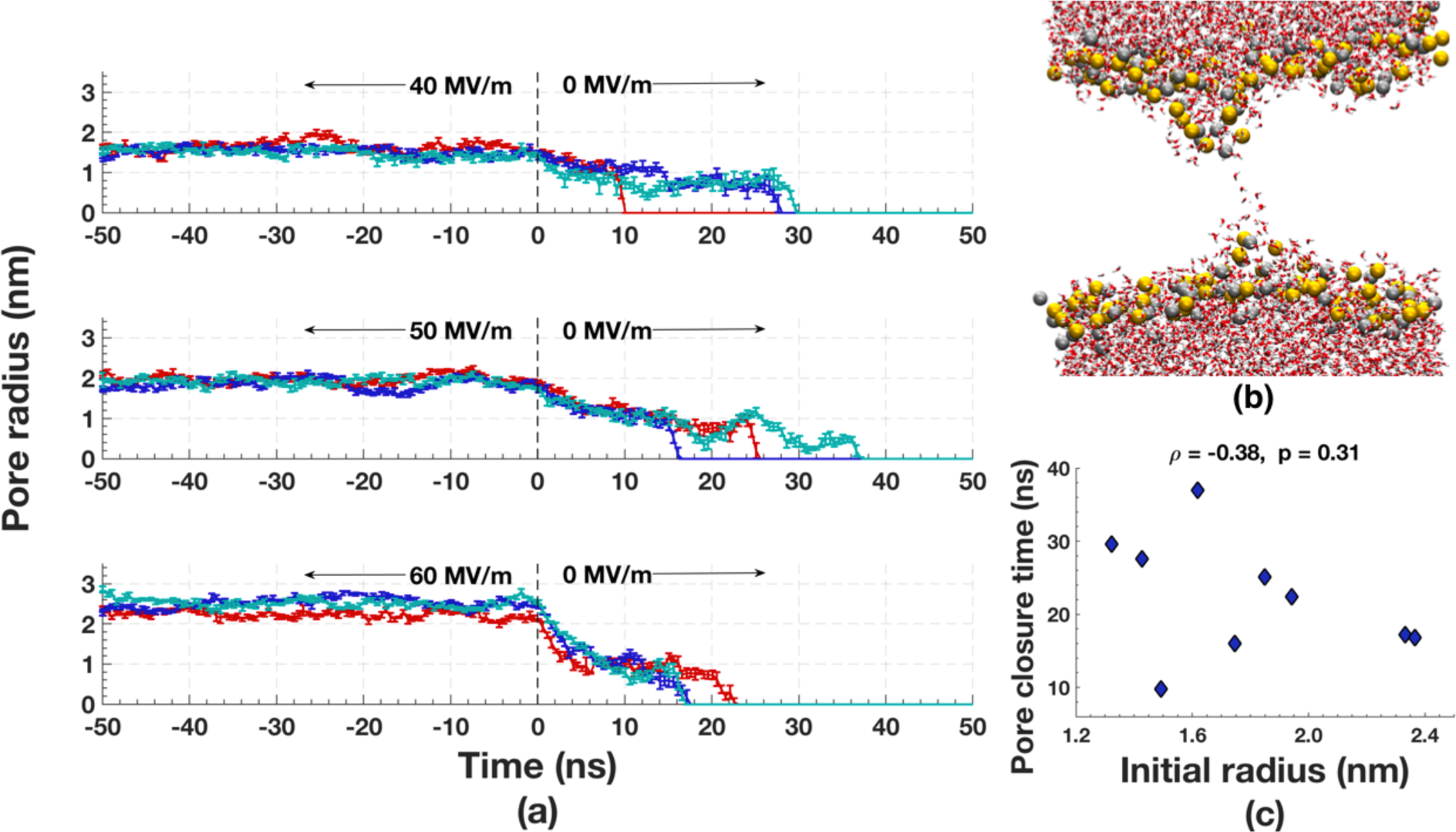
(a) Kinetics of pore annihilation for three different pore sizes. Three independent simulations indicated with different colors (red, blue, cyan) for each pore size are shown. Error bars indicate standard deviation of radius (Supplementary Material). (b) Snapshot of a typical lipid electropore just before final collapse of the membrane-spanning water column. (Red and white: water O and H; gold and silver: P and N of lipid head groups; lipid tails hidden for clarity). The pore is shown 9.5 ns after removal of 40 MV/m sustaining field, 0.5 ns before complete closure (simulation plotted in red in panel a, top plot). (c) Scatter plot showing a lack of correlation between the initial pore radius (mean of the first 1 ns after the electric field removal) and the pore closure time.

In conclusion, our experimental data indicate that lipid electropores in GUVs are created within a few nanoseconds and that most are annihilated within a few tens of nanoseconds, consistent with molecular simulations, but in contrast with the typical persistent electropermeabilization (many seconds to minutes) observed in living cells. Furthermore, the magnitudes of the increases in intravesicular dye concentration are much smaller with nanosecond pulsed electric fields than those observed in the presence of the pore-forming peptide melittin or exposure to influenza virus at low pH. ^36^ Nanosecond bipolar pulse cancellation,^28^ a phenomenon recently described in cells, was not observed in GUVs. The absence of persistent electropermeabilization and nanosecond bipolar cancellation of GUVs suggests that the electropermeabilization of cells involves structures and processes that go beyond transport through lipid pores, and that models of electroporation must be modified accordingly.

## Acknowledgements

This work was supported in part by the Division of Intramural Research of the NICHD. EBS and PTV were supported by AFOSR MURI grant FA9550-15-1-0517 on “Nanoelectropulse-Induced Electromechanical Signaling and Control of Biological Systems”, administered through Old Dominion University.

## Supporting Information

Details of methods and additional data analyses are provided in the supplementary documentation.

