## Supplementary Material for "Ultra-fast electroporation of giant unilamellar vesicles — Experimental validation of a molecular model"

#### Molecular Simulations

***Simulation conditions.*** For molecular dynamics (MD) simulations we used GROMACS 4.6.6. **(van der Spoel *et al.* 2005; van der Spoel *et al.* 2014)**, on the Old Dominion University High Performance Computing cluster (http://www.odu.edu/hpc) with CHARMM36 lipid topologies and force field parameters (http://mackerell.umaryland.edu/charmm_ff.shtml) **(Klauda *et al.* 2010)** and the TIP3P water model **(Jorgensen *et al.* 1983).** The online tool *CHARMM-GUI: Membrane Builder* was used to create lipid bilayer systems **(Jo *et al.* 2008)**. All simulations were carried out under the NPT ensemble with a 2 fs time step. Systems were coupled to a temperature bath at 310 K using the velocity-rescale coupling algorithm **(Bussi *et al.* 2007)**, and to a pressure bath at 1 bar using the Berendsen algorithm **(Berendsen *et al.* 1984)** with a relaxation time of 1 ps and compressibility of $4.5 \times 10^{-5}$ $bar^{-1}$ semi-isotropically applied in the normal and in-plane directions relative to the membrane. Bond lengths were constrained using the LINCS algorithm **(Hess *et al.* 1997)** for lipids and SETTLE **(Miyamoto and Kollman 1992)** for water. Short-range electrostatic and Lennard-Jones interactions were cut off at 1.0 nm. Long-range electrostatics were calculated by the PME algorithm **(Essmann *et al.* 1995)** with conductive periodic boundary conditions.

***Systems and structures***. To lipid bilayer systems containing 128 POPC (1-palmitoyl-2-oleoyl-*sn*-glycero-3-phosphatidycholine), 64 in each leaflet, and approximately 12,000 water, we added 22 $K^+$ and 22 $Cl^-$ using the GROMACS function “genion”. Box dimensions are approximately 7 nm × 7 nm × 11 nm, and the KCl concentration is about 125 mM (based on a volume that excludes the membrane interior and interface where water density is less than 90% of the bulk water density).

The system was equilibrated for 100 ns until a constant area per lipid was achieved (**Vernier *et al.* 2009)**. A porating electric field of 250 MV/m was applied along the z-axis (normal to the bilayer plane) following the method used in **(Ziegler and Vernier 2008)** to induce formation of an electropore through the lipid bilayer. A procedure similar to the one described in (**Fernández *et al.* 2012)** was used to produce stable electropores at different stabilizing field amplitudes: 40, 50 and 60 MV/m. Three independent trials were carried out for each stabilizing field value by randomizing velocities after pore formation (GROMACS MDP file: gen_vel = yes and gen-seed = -1). After 100 ns of stabilization, the electric field was reduced to zero, and the simulations were run for another 100 ns.

***Pore radius and pore closure time.*** Pore geometry was extracted from GROMACS structure (gro) files generated from the simulations every 100 ps and analyzed with custom MATLAB scripts. Pores were centered in each frame using POPC density files generated with the GROMACS command "g_density". The center of the pore was determined as follows: (1) POPC density versus spatial axes plots are smoothed using a Butterworth low-pass filter generated by the native MATLAB function "butter"; (2) The region containing the pore is defined by decreases in POPC density in x, y, and z. The dip defining the pore is chosen as the region in the density profile that is less than 34% of the full range of densities. The pore center, which is in the middle of this dip region, is placed in the center of the simulation box, translating all atoms in the frame accordingly. After centering the pore, the box is divided into 0.2 nm slices in the z-direction. Water molecules in each slice are located, and outliers in the x- and y-directions are eliminated using the native MATLAB function "isoutlier", which defines an outlier as a value that is more than three scaled median absolute deviations away from the median. These outliers represent occasional random water intrusions into the lipid bilayer far from the electropore. Next, the center of each z-slice in x and y is determined using the mean of water oxygen locations, and the distance in the x:y plane of each water molecule (oxygen atom) from the slice center is calculated. The ***water-based radius*** of each slice is defined as the mean plus two standard deviations of this set of distances. This generates a water-column radius versus z-slice location vector for each frame. The pore radius ($r_{pore}$) in a frame is defined to be the mean water-column radius in the 1 nm-long region around the z-center of the pore. Pore radius versus time is plotted in Figure 3a using the mean pore radius of every 5 frames (500 ps). Error bars show the standard deviation of the pore radius value for the same 5 frames.

Pore closure time is defined as the time of the first of three consecutive frames with $r_{pore} = 0$; that is, three consecutive frames in which 5, 0.2 nm thick z-slices around the box center are water-free.

### Experimental Methods

***Giant Unilamellar Vesicles***. Pure 1-palmitoyl-2-oleoyl-*sn*-glycero-3-phosphocholine (POPC, Avanti Polar Lipids) GUVs containing 200 mM sucrose dissolved in PIPES (1 mM EDTA, 1 mM HEDTA, 10 mM PIPES, 100 mM KCl, pH 7.4) were prepared with a gel-assist protocol (**Weinberger *et al.* 2013**, **Stein *et al.* 2017, Haldar *et al.* 2019**) and stained with DiD (Invitrogen) **(Haldar *et al.* 2019)**. GUVs were pipetted 15 minutes before the recording into cover glass chambers (Thermo Scientific™ Nunc™ Lab-Tek™ II) filled with 200 μM calcein (MP Biomedicals) and 200 mM glucose in PIPES solution where parallel tungsten wire electrodes were pre-positioned in the chamber well.

***Microscopy***. Images were acquired on a Zeiss LSM 880 confocal microscope using a 63×, oil immersion, 1.4 NA objective. The size of the confocal pinhole was set to one Airy disk at 633 nm. Calcein and DiD were excited with 488 nm and 633 nm lasers respectively, by alternating every 0.03 ms lines during raster image formation, and fluorescence was measured (emission bands $\lambda_{calcein}$: 500–561 nm and $\lambda_{DiD}$: 635–735 nm) at a rate of one frame per second using separate aquisitions (tracks) to minimize crosstalk.

***Pulsed electric field exposure.*** 6 ns or 2 ns – full-width at half-maximum (FWHM) – electric pulses from FID GmbH pulse generators (6 ns: FPG 10-10NK, 2 ns: FPG 10-1CN6V2, Burbach, Germany) were delivered to lipid vesicles via parallel tungsten wire electrodes 80 μm apart **(Sözer *et al*. 2017, Sözer and Vernier 2019)**. Pulse exposures were either unipolar or bipolar with equal positive and negative phase amplitudes and were monitored and recorded with an oscilloscope. Typical waveforms are shown in Figure S1. Electric field amplitudes at the vesicles were computed using COMSOL multiphysics (COMSOL AB, Stockholm, Sweden). For the 6 ns pulse responses in Figure 1, 10, 35 MV/m pulses were delivered at 1 kHz; for the 2 ns pulse responses in Figure 2, 40, 45 MV/m pulses were delivered at 1 kHz.

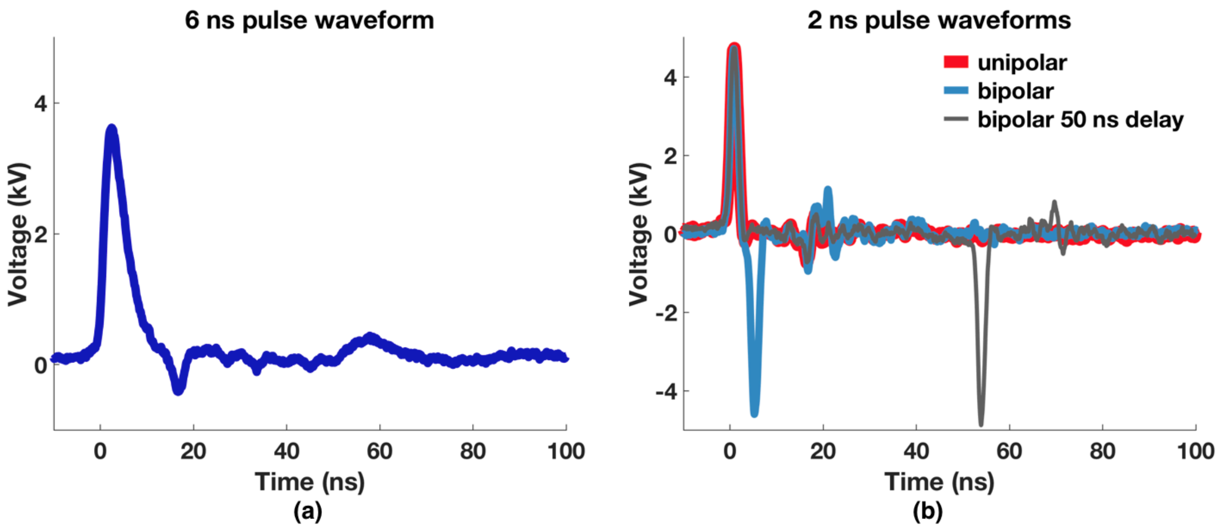


*Figure S1. Typical electric pulse waveforms recorded during the experiments. (a) 6 ns exposure. (b) 2 ns exposures: unipolar (red), bipolar without any interphase time interval (blue), and bipolar with 50 ns between the phases (gray).*

### Vesicle size correlation with fractional fluorescence change

Bioelectromagnetic theory predicts that a GUV (a spherical dielectric shell) in a uniform electric field in a conductive medium will develop a peak induced membrane potential proportional to its radius **(Pauly and Schwan 1959)**.

$$\Delta\psi_m = \frac{3}{2} E_0 r\,(1 - e^{-t/\tau_m}) \qquad \text{(S1)}$$

where $\psi_m$ is the membrane potential, $t$ is the duration of the electric field exposure, and $\tau_m$ is the membrane charging time constant. However, the positive correlation between vesicle radius and calcein fluorescence increase predicted by classical electroporation theory is not seen in our data (Figure S2), in contrast to experiments where longer pulse durations were used **(Mauroy *et al.* 2012)**. This size-independence of membrane permeabilization is also seen in cells, when the pulse duration is much shorter than $\tau_m$ (approximately 100 ns in the current set of experiments). Mathematically, this comes from the linear approximation of the exponential term in equation S1 for small $t/\tau_m$ , since $\tau_m$ is directly proportional to vesicle size **(Sözer *et al.* 2017, Stewart *et al.* 2004)**. The negative correlation of vesicle size and fractional fluorescence change shown in Figure S2a is not significant when a smaller time range before and after pulse delivery is used to calculate fractional fluorescence change (Figure S2b and c). One interpretation is that a rapidly opening and closing pore population forms independent of vesicle size, causing an immediate fluorescence change at pulse delivery, and that the formation of longer-lasting (seconds) pores is negatively correlated with vesicle size. The reason for this dependence needs further investigation with a larger sample size.

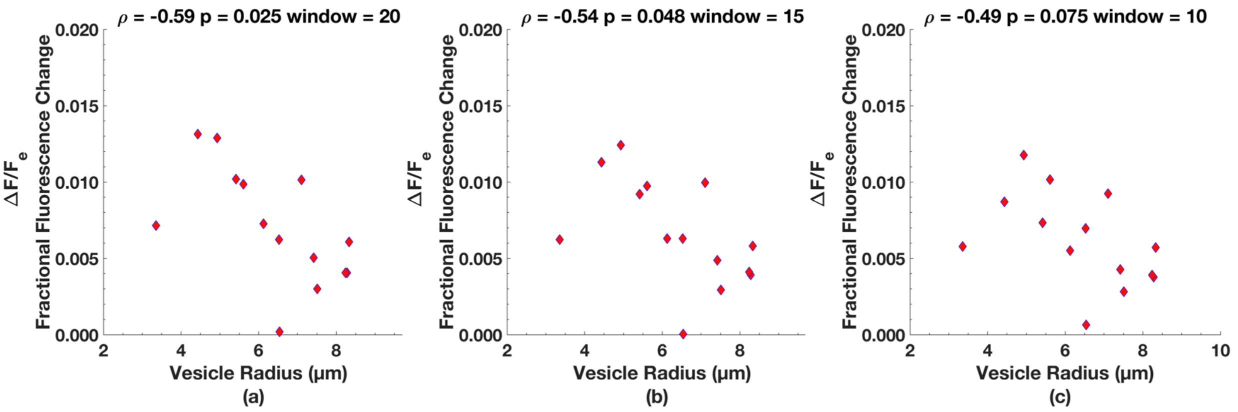


*Figure S2 Vesicle size dependence of fractional fluorescence change calculated using window size (a)20 (b) 15 and (c) 10 around first pulse train delivery.*

### Kinetics of fluorescence change and electrodiffusion of calcein

From the kinetics presented in Figure 1b, we wonder if *6 ns (unipolar)* pulses form some longer lifetime pores (seconds) in the membrane in addition to a much larger number of fast-closing (tens of nanoseconds) pores. Longer lifetime pores can also explain the slower rise time of the responses to 2 ns bipolar pulses without any interpulse delay, which induces a high transmembrane potential across the membrane for 4-5 ns, resulting in the formation of a population of longer-lifetime pores similar to those produced by the unipolar 6 ns pulse exposures. Lipid pores with longer lifetimes are observed experimentally and in simulations when a low membrane potential is present **(Abidor**

***et al.* 1979, Sengel and Wallace 2016, Fernandez *et al.* 2012).** This hypothesis suggests that longer duration pulses are more likely to form longer lifetime pores. See Figure S3 overlays of 6 ns unipolar and 2 ns bipolar waveforms and calcein uptake kinetics. A small population of longer lifetime pores with bipolar pulses without any interpulse delay is also consistent with MD simulations in which the electric field direction is reversed after pore formation. Field reversal did not affect pore size or lifetime. The pore continues to evolve as if the change in the direction of the field did not happen. The longer lifetime pore population could also lead to a change in the system that affects response to subsequent pulse trains.

In contrast, with *2 ns unipolar* pulses in Figure 2 (either single or with 50 ns interpulse delay), the contribution of longer lifetime pores in the pore population is lower, thus we see sharp increases (faster than our recording speed of 1 fps) in calcein fluorescence intensity.

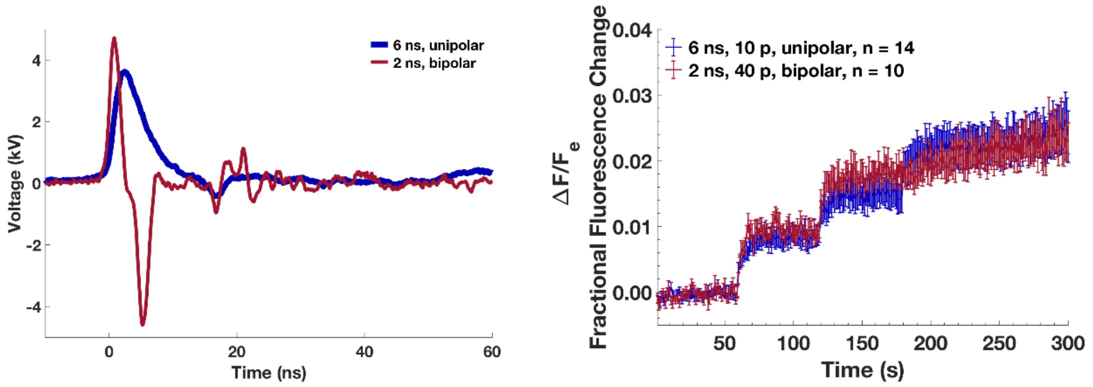


*Figure S3. Left: overlay of 2 ns bipolar pulse with no interpulse delay and 6 ns unipolar pulse. Right: calcein uptake kinetics after exposure to two different pulse trains, consisting of the pulses shown on the left panel. Maroon: a train of 40 bipolar pulses (45 MV/m, 2 ns, 1 kHz repetition rate) were delivered at 60 and 120 seconds into the recording (also shown in the main manuscript figure 2). Blue: a train of ten unipolar pulses (35 MV/m, 6 ns, 1 kHz repetition rate) delivered at 60, 120 and 180 seconds into the recording (also shown in the main manuscript figure 1b).*

The kinetic changes following pulse exposure to a train of ten pulses of 6 ns duration and field strength of 35 MV/m (Figure 1b) can be modeled as an "instantaneous" jump (unresolvable at 1 Hz imaging) at the start of a pulse sequence followed by a kinetic term with a single time constant. For the data presented in Figure 1b, both piecewise and global fitting support the hypothesis that the instantaneous jump, the contribution of the kinetic term, and the time constant are the same. The basic model is

$$IJ + A\,(1 - \exp\,(-t/\tau)) \qquad \text{(S2)}$$

where $IJ$ is the "instantaneous" change measured in the first image frame, $A$ is the amplitude associated with the kinetics resolved in subsequent image frames, and $\tau$ is the time constant. Table S1 summarizes the fitted data obtained using three parameter, piece-wise fitting, after subtracting the starting plateau value, and Figure S4 shows the data with the fit. Regression of all parameter values, as a function of pulse sequence also indicates no linear dependence (slope not statistically different from 0) supporting the hypothesis that each pulse sequence, on average, is an independent realization of the underlying creation of pores with approximately half of the signal change due to electrophoretic transport (drift) through short lived pores and the other half due to diffusive transport through longer lived pores.

*Table S1Three parameter piece-wise fitting of kinetics of fluorescence change after 6 ns, 35 MV/m pulse train (Fig 1b main manuscript) to equation S2*

| | Pulse Train | | |
|---|---|---|---|
| *ΔF/Fe* | 1 | 2 | 3 |
| Amplitude (*A*) | 0.005 | 0.003 | 0.004 |
| Instantaneous Jump *(IJ)* | 0.004 | 0.004 | 0.004 |
| Total Change | 0.009 | 0.007 | 0.007 |
| 95% Confidence | 0.002 | 0.002 | 0.001 |
| $\tau$ (s) | 7.7 | 9.8 | 12.0 |
| 95% Confidence | 3.5 | 7.3 | 5.2 |

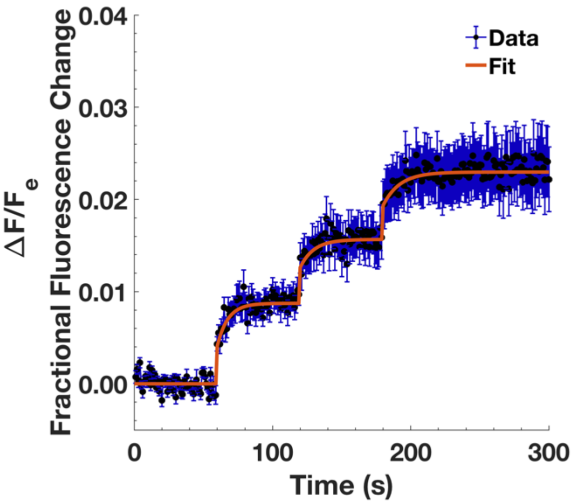


*Figure S4. Three parameter piece-wise fitting of data from main manuscript Figure 1b where GUVs exposed to three 1 kHz trains of ten 6 ns electric pulses at field strengths of 35 MV/m, delivered 1, 2, and 3 minutes into the recording (n = 13).*

We next consider whether the ~1% changes in fluorescence observed using 6 and 2 ns pulses are consistent with a population of transient pores of nanosecond duration undergoing electrodiffusion. Pore-mediated transport of calcein was modeled using a simple electro-diffusive transport model as described previously **(Sözer *et al.* 2018)**. Briefly, following the Nernst-Planck formalism $J_{p,}$ electrodiffusive transport through a single cylindrical pore was defined as sum of two components, the diffusion term ($J_{diffusion}$) and the drift term ($J_{drift}$):

$$J_p = J_{\text{diffusion}} + J_{\text{drift}} \quad \text{(S3)}$$

$$J_{\text{diffusion}} = \frac{\pi\, r_{\text{pore}}{}^2 D_S\, c}{l_{\text{pore}} + \frac{\pi\, r_{\text{pore}}}{2}} \quad \text{(S4)}$$

where $D_s$ is the diffusion coefficient, $r_{pore}$ is the pore radius, $l_{pore}$ is the length of the pore (4 nm in these calculations), and $c$ is the concentration difference from one side of the membrane to the other. The drift term is

$$J_{\text{drift}} = \frac{1}{2} \frac{\pi\, r_{\text{pore}}{}^2 D_S\, c}{l_{\text{pore}}} \frac{q_{\text{e}} z V_m}{kT} \quad \text{(S5)}$$

where $V_m$ is the transmembrane potential, which can be estimated with equation S1 using the electric field amplitude and $r_{vesicle}$ vesicle radius , which is 6 µm on average in our experiments.

We assume that following electroporation, 1/1000 of the vesicle's surface is populated by 1 nm pores ($n_{pore,fast}$ = 1.4e5) with a pore formation time of 1.2 ns, after which drift and diffusion processes begin. The drift lasts only as long as the duration of the pulse while diffusion continues for 50 ns . With these assumptions we get the values in Table S2.

*Table S2 Calculations of electrodiffusion of calcein through short-lived pores*

| | 6 ns, 35 MV/m, 10 p | 2 ns, 45 MV/m, 40 p |
|---|---|---|
| $V_m$ (V) | 300 | 400 |
| $J_{drift}$ (mol/s) | 3.2e-16 | 4.3e-16 |
| $J_{diffusion}$ (mol/s) | 1.1e-20 | 1.1e-20 |
| $n_{pore,\ fast}$ | 1.4e5 | 1.4e5 |
| $c_{intravesicular,drift,\ single\ pulse}$ (M) | 2.5e-7 | 5.4e-8 |
| $c_{intravesicular,diffusion,\ single\ pulse}$ (M) | 1e-10 | 9e-11 |
| $c_{intravesicular,\ pulse\ train}$ (M) | 2.4e-6 | 2.1e-6 |
| $\Delta F/F_e$ | 0.01 | 0.01 |

These calculations show that the transport of calcein we observed as an instantaneous jump following the pulse sequence can be explained by transport through electropores with ~50 ns lifetime even when we limit the number of pores to a small fraction of the vesicle surface. The slower kinetics (seconds) observed with 6 ns unipolar pulses are attributed to a much smaller number of longer lived pores undergoing diffusive transport alone.

For example, in the 6 ns pulse case if we replace half of the short-lived pores with only ten pores with a lifetime of 1 second at full transport capacity, we get approximately equal contribution of short and longer lifetime pores as listed in Table S3.

*Table S3 Calculations of electrodiffusion of calcein through short-lived and longer-lived pores*

| | 6 ns, 35 MV/m, 10 p |
|---|---|
| $V_m$ (V) | 300 |
| $J_{drift}$ (mol/s) | 3.2e-16 |
| $J_{diffusion}$ (mol/s) | 1.1e-20 |
| $n_{pore, fast}$ | 7e4 |
| $n_{pore, slow}$ | 10 |
| $c_{intravesicular,drift, single pulse}$ (M) | 1.2e-7 |
| $c_{intravesicular,diffusion, single pulse}$ (M) | 1.3e-7 |
| $c_{intravesicular, pulse train}$ (M) | 2.5e-6 |
| $\Delta F/F_e$ | 0.01 |
